# Open Hardware: Towards a Fully-Wireless Sub-Cranial Neuro-Implant for Measuring Electrocorticography Signals

**DOI:** 10.1101/036855

**Authors:** David Rotermund, Jonas Pistor, Janpeter Hoeffmann, Tim Schellenberg, Dmitriy Boll, Elena Tolstosheeva, Dieter Gauck, Heiko Stemmann, Dagmar Peters-Drolshagen, Andreas K. Kreiter, Martin Schneider, Steffen Paul, Walter Lang, Klaus R. Pawelzik

## Abstract

Implantable neuronal interfaces to the brain are an important keystone for future medical applications. However, entering this field of research is difficult since such an implant requires components from many different areas of technology. Since the complete avoidance of wires is important due to the risk of infections and other long-term problems, means for wireless transmitting data and energy are a necessity which adds to the requirements. In recent literature many high-tech components for such implants are presented with remarkable properties. However, these components are typically not freely available for your system. Every group needs to re-develop their own solution. This raises the question if it is possible to create a reusable design for an implant and its external base-station, such that it allows other groups to use it as a starting point. In this article we try to answer this question by presenting a design based exclusively on commercial off-the-shelf components and studying the properties of the resulting system. Following this idea, we present a fully wireless neuronal implant for simultaneously measuring electrocorticography signals at 128 locations from the surface of the brain. All design files are available as open source.

## Introduction

There is nothing more drastic in a person's life than losing motor control over the own body (e.g. by a neuro-degenerative disease, stroke or paraplegia), getting blind or losing limbs. The actual state of medical technology has only limited options for helping this group of people. Results from brain-research suggest that it should be possible to build technical medical devices which interact with the neuronal activity patterns of the brain to ease the loss of life quality and partially restore functionality (e.g. creating visual perception [1, 2, 3] and extracting information from neuronal activities [4, 5, 6, 7]). Even with the limited knowledge of today, astonishing assisting systems for this group of people are possible [8, 9, 10, 2]. One important example are invasive brain-computer interfaces (BCI), which allow to control computers or robot-arms by evaluating the actual spatiotemporal cortical activity patterns (e.g. [11, 4, 6, 5, 12, 8, 9, 13, 14, 15] and many more).

Transferring such systems into the daily medical routine remains a highly challenging task. Effective control of external devices with invasive BCIs requires recording of neuronal data with high temporal and spatial resolution, which is best achieved with intracortical recordings. However, intracortical implantation of electrodes might lead to brain damage. Furthermore, recording quality usually degrades over time due to formation of scar tissue around the electrodes. An applicable compromise are electrocorticography (ECoG) signals, recorded from the surface of the brain or the dura mater, which still contain detailed information usable for BCI [16, 17, 18, 19, 20, 21]. Further requirements for an implantable interface are long-term stability (up to several decades), bio-compatibility, and persistence against humidity. From a functional point of view these systems need to provide a high spatial and temporal resolution to measure and/or change the neuronal activity patterns in the human brain.

To achieve the functionality of neuro-prostheses, complex data analysis procedures need to be applied in real time [16, 22, 5, 23, 24]. With current technology, it is not possible to perform this computationally extensive data processing with processor units placed inside the human body. The main reason is the high amount of heat produced by the processors, which would lead to tissue damage [25, 26, 27]. Therefore, a neuro-prosthesis needs to consist of two parts: an implanted device with recording (and optionally stimulation) capabilities and an external analysis/ control system. However, a tethered data transmission between the two parts has an inherent risk of infection [28, 29, 30], cerebral fluid loss, as well as bio-mechanical problems in chronic applications. A solution of this problem is a wireless connection between the implant and the base station. This would allow to fully embed the implant inside the human body without physical connections. It would be even more advantageous if the implant could completely be placed inside the human skull (e.g. for keep the fluidic environment around the brain intact). Following these requirements, the implant has to be capable of exchanging data wirelessly with an external base station through skull, fat, fluids and skin. In order to avoid components with limited lifetimes like batteries, a wireless power link has to provide the energy for the implant.

Although there are approaches for wireless data exchange [31] (using various technologies, e.g. Ultra-Wide Band [32], Offset Quadrature Phase Shift Keying [33], Amplitude Shift Keying [34], Frequency Shift Key Modulation [35], RF backscattering [36] and RF ID Technology [37]) and energy transmission systems [38, 39, 40], especially for neuronal implants, these systems are not available on the market. Thus, it is nearly impossible for other groups to obtain and (re)use these systems. We would like to support other researchers in the field by providing them with a wireless energy and data interface, and thus push their research and enable them to focus on other functional parts of their system. Therefore, we present here a neuro-implant for sub-cranial implantation that is based on commercial off-the-shelf (COTS) components.

Our system is targeted for animal experiments. In the long run, we aim at a system for human medical applications. In 2010 we started to design an application specific integrated circuit (ASIC) [41] which was able to communicate with the undocumented analog-digital-converter on the Intan Tech bio-signal amplifier RHA 2116 for collecting electrophysiological data. Later we improved this ASIC design by upgrading it with support for the wireless module ([42, 43]) which we present here in detail. Those first prototypes were based on large and fully rigid FR-4 PCBs. Because it is important that the implant follows the curvature of the brain [44], we started preparation for integrating all components on an industry grade flexible PCB-foil.

Here we present our wireless module and its base station for exchanging data and providing energy to the implant. For the first time, we make all design files (circuit diagrams, board designs, test boards, firmware and software) available as open source. Furthermore, we re-implemented the functionality of our ASIC as a firmware for a Microsemi IGLOO nano FPGA. We also wrote a second IGLOO nano FPGA firmware for supporting the newer Intan RHD2132 with better ADC performance, instead of using the undocumented ADC features of the Intan RHA2116. Both firmwares are also part of the open-source package as well as test boards for the Microsemi nano FPGA and the Intan RHD2132. This allows us to present a design which can be built completely from commercial off-the-shelf components and make it available as open-source. Since the Microsemi nano FPGA and the Intan RHA/ RHD are all available as bare dies, the size of the system is suitable for an implant usable for human medical applications.

In parallel, we investigated how our ASIC with our open-source wireless module and Intan RHA2116 operates on an industry grade flexible PCB-foil. We analyze the results and report which problems arose. Furthermore, we are examining the temperature distribution around the implant in measurements and simulations.

## Results

### System concept

Our design goal was to build a system that can be implanted completely subcranially, which is supplied with energy via a wireless link (without any implanted batteries) and which exchanges data wirelessly with an external base station. Figure 1 shows the functional blocks necessary for such a system.

**Figure 1:**
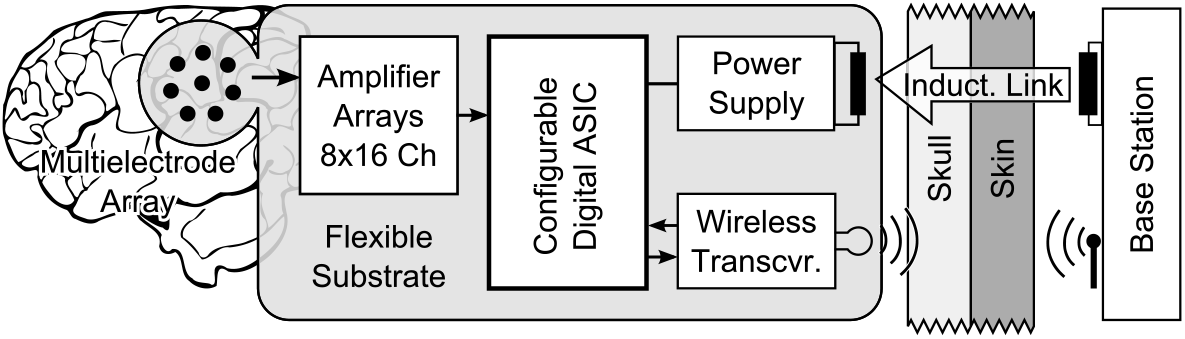
Concept of the implant with its base station.

An array of electrodes serves as an interface between the brain tissue and a set of integrated analog signal amplifiers with band-pass properties. After the amplification of the neuronal signals, analog-digital converters digitize the incoming signals and generate several digital data streams. An Application Specific Integrated Circuit (ASIC) filters these data streams according to user-defined specifications and merges the parallel streams into a single one, optimized for a minimal bandwidth. The condensed data stream is re-packed into transmission packages and transmitted via an RF transceiver data link to an external base station. The base station receives the data packages and unpacks, checks, and repacks them. These newly built packages, optimized for fast processing by 32 or 64-bit CPUs, are sent via Ethernet to an external PC for further use (e.g. visualization and analysis). From the external PC, the base station receives instructions about the user defined parameters for the data processing of the implant and transmits them to the implant to set the desired configuration in the ASIC. Beside the bi-directional wireless data exchange, the implant collects energy from an inductive wireless power link for power supply.

### The wireless module

The presented wireless module incorporates two connected sub-segments: One which supplies the implant wirelessly with energy and the other one for wireless communication. Figure 2 visualizes all the necessary components plus its external counterparts. Figure 3 shows a PCB realization of that block diagram with a total size of 20 mm × 20 mm × 1.6 mm. In the following section, both functional blocks are described in detail.

**Figure 2:**
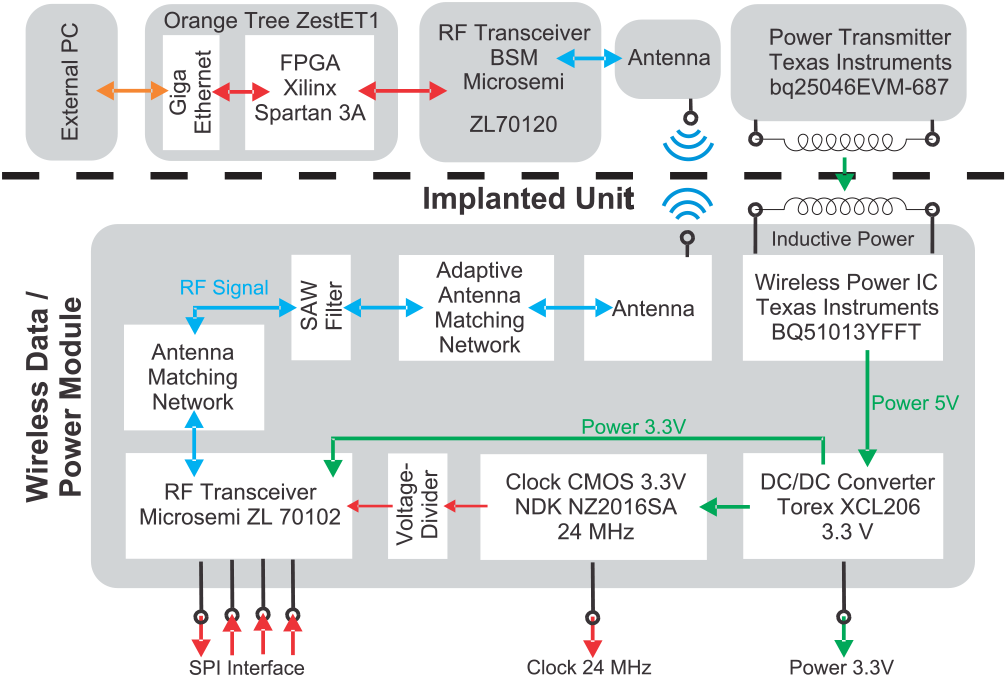
Overview of the components required for realizing the presented system concept of the wireless energy and data link.

**Figure 3:**
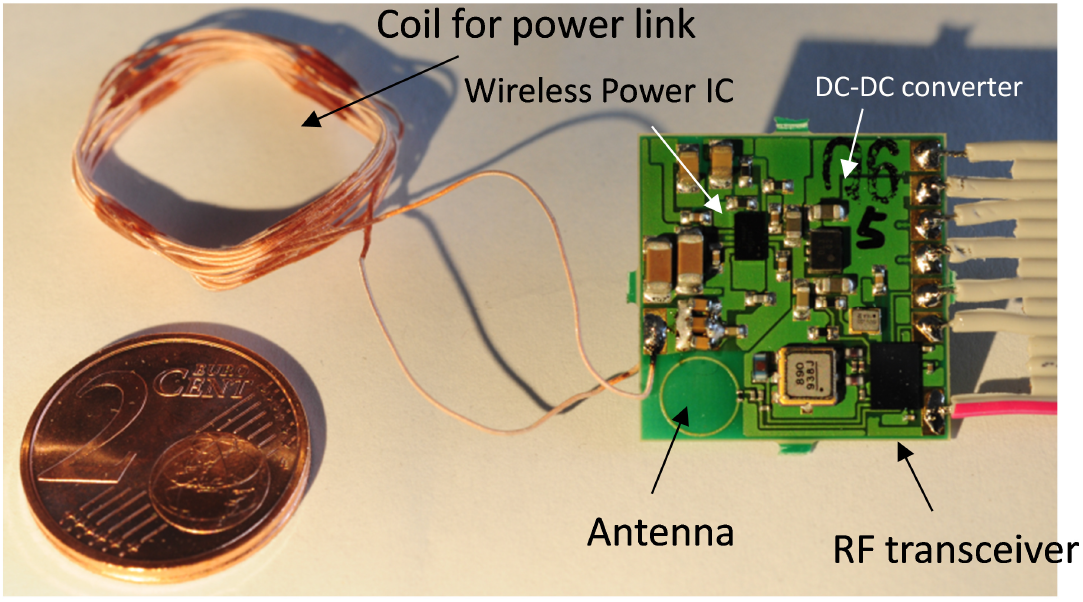
Realization of the wireless data / power module on a 0.15mm thick FR4 board (20×20×1.6mm3) with its hand wound coil for the inductive power link.

**The power supply:** The concept of the wireless power link was designed based on the Texas Instruments (TI) bqTESLA system [45]. TI designed these products for wirelessly recharging mobile devices, e.g. MP3-players and smart-phones, based on the QI standard. Consequently, the corresponding components - designed for an integration into a mobile device - are highly miniaturized and designed for high efficiency. In theory, this link can deliver up to 5 Watt [46]. Between the energy receiver and transmitter an ongoing communication regulates the properties of the wireless energy link dynamically and load-dependently. The frequency of the power link is dynamically regulated between 110 kHz and 205 kHz [46] and depends on the amount of the consumed power on the secondary side (implant). This regulation, in combination with a fixed resonance frequency of the receiver, helps to prevent harvesting too much energy on the implant site, that would lead to unnecessarily heating up the implant as well as the surrounding tissue.

On the primary side (base station) we used the bqTESLA wireless power evaluation kit (bq25046EVM-687) as a off-the-shelf low-cost power transmitter [46]. On the secondary side, a BQ51013YFFT IC as power receiver is part of the design [47]. This chip-sized ball grid array (BGA) contains the means to communicate with the external energy transmitter, rectification of the inducted AC wave, and voltage regulation. The receiver IC delivers a 5V power rail. If this IC is used, then two important design aspects have to be considered: 1.) Many of the required ceramic capacitors on the side of the secondary coil need to be rated for 50V. As a result, the capacitors with larger capacitance are too thick for some target areas of implantation. Therefore, it is necessary to split them into several smaller parallel capacitors. 2.) Due to the small distance between the balls of the BGA, it was not possible to contact important pads in a typical fashion. Thus, it is necessary to place via holes underneath the pads for the BGA package. This requires the via to be filled up and closed with a planar surface, which is quite demanding for the (external) manufacturer of the printed circuit board (PCB).

The 5V output of the power receiver IC is too high for operating the RF transceiver and other active components. Thus, a highly efficient and very small DC/DC converter is required. We applied a Torex XCL 206 step-down micro DC/DC converter with built-in inductor which only requires two small capacitors as external components [48]. In the expected operating point, it works with an efficiency over 80%. Due to its switching nature, PI filters, for smoothing the DC supply rail, are advised on the consumer side. However, additional capacitive loads exceeding 50*μ*F, by e.g. PI filters and block capacitors for the ICs, cause problems and loads beyond 70*μ*F stopped the DC/DC converter from working at all.

**Data Transfer:** The wireless data transfer is based on Microsemi ZL70102 transceivers [49]. The RF transceiver operates in the Medical Implant Communication Service frequency band (MICS, 401 - 406 MHz) and is commercially available for medical applications including implants. The transceiver establishes a bi-directional wireless link, using 4-FSK or 2-FSK mode of operation. In order to achieve a high data rate, especially for a continuous data streaming, it is necessary to provide a large extra memory for the controller, which is operating the transceiver via SPI on the implant site. This is necessary because pauses of unknown origin in the data transfer of up to 80 ms may randomly occur. The transceiver is available as a chip-sized BGA.

The ZL70102 requires several external components. Among those is a 24 MHz clock. We used a very small (2 mm × 1.6 mm × 0.7 mm) CMOS 3.3 V clock from NDK (NZ2016SA) [50]. Besides driving the transceiver, it also provides a clock signal for other components (e.g. ADCs, microcontroller, FPGAs or ASICs (e.g. [51]) for data processing).

Between the RF transceiver and the antenna (circular loop antenna with 5mm diameter), we installed an adaptive antenna-matching circuit with a SAW filter (RF Monolithics RF3607D, 403.5 MHz SAW filter) [52]. The adaption of the matching circuit is accomplished by using two tunable capacitors which are part of the ZL70102. Those are optimized by the transceiver automatically. The SAW filter is one of the largest components (3.8 mm × 3.8 mm × 1.0 mm) on the implant.

For the base-station module outside the body, a solution based on a Microsemi ZL70120 was designed [53]. Among other components, this Microsemi RF transceiver base station module contains a ZL70102, antenna-matching circuit and a clock. We designed a simple PCB for this module and added a 50 Ohm rectangular loop antenna to it (see Figure 4). Via SPI, we operated this transceiver base station module with a FPGA using a custom firmware. This FPGA is part of a board (Orange Tree ZestET1) with Gigabit Ethernet connectivity [54]. This allows to stream the data from the implant to an external PC via TCP/IP. The base station also supplies the implant with control sequences from the PC using the other direction of communication.

**Figure 4:**
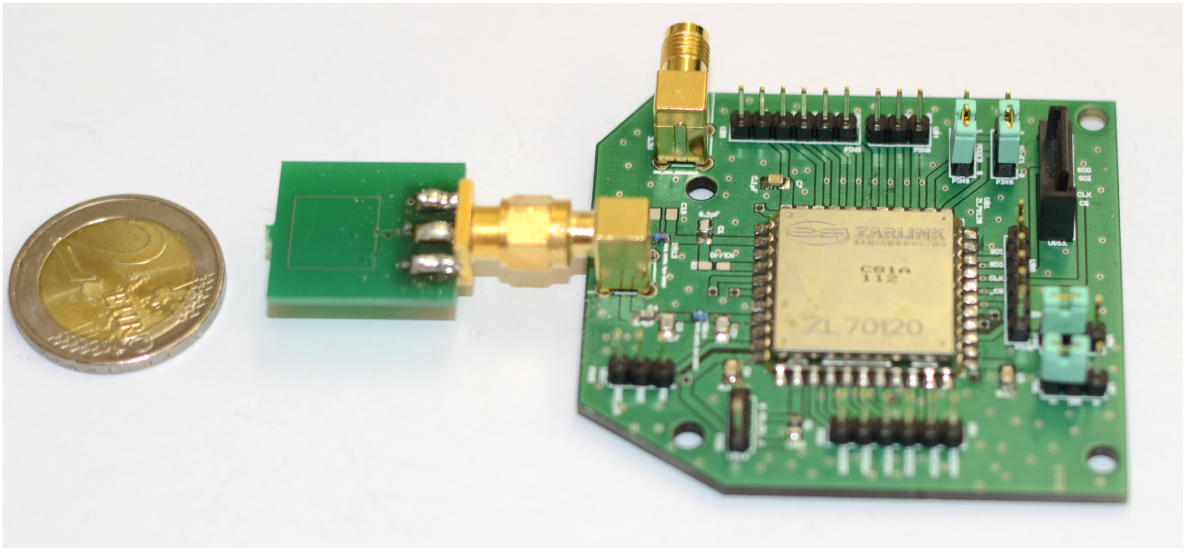
Base station RF transceiver module based on a Microsemi ZL70120 module and an in-house designed antenna. The rest of the base station has to be connected via SPI to this module.

### The implant prototype

The described system for the implant was realized (see Figure 5) with 128 gold electrodes in an area of 9mm × 17mm with a diameter of 0.4mm for the individual electrodes and a center-to-center electrode distance of 1.4mm on a flexible 50*μ*m thick PCB-foil (DuPont Pyralux AP). This PCB-foil has a size of 34mm × 79mm. It can be folded at 3 lines (see Figure 6) to further reduce the overall size of the implant as shown in Figure 7. All electrical components are placed on one side of the two-layered PCB-foil with respect to the polyimide process developed for future implementations. The weight of the implant is 1.72g, and it fits into a volume of 4mm×24mm×32mm excluding the power-link coil. The coil has a square shape with a side length of 22mm and a thickness of 2.2mm.

**Figure 5:**
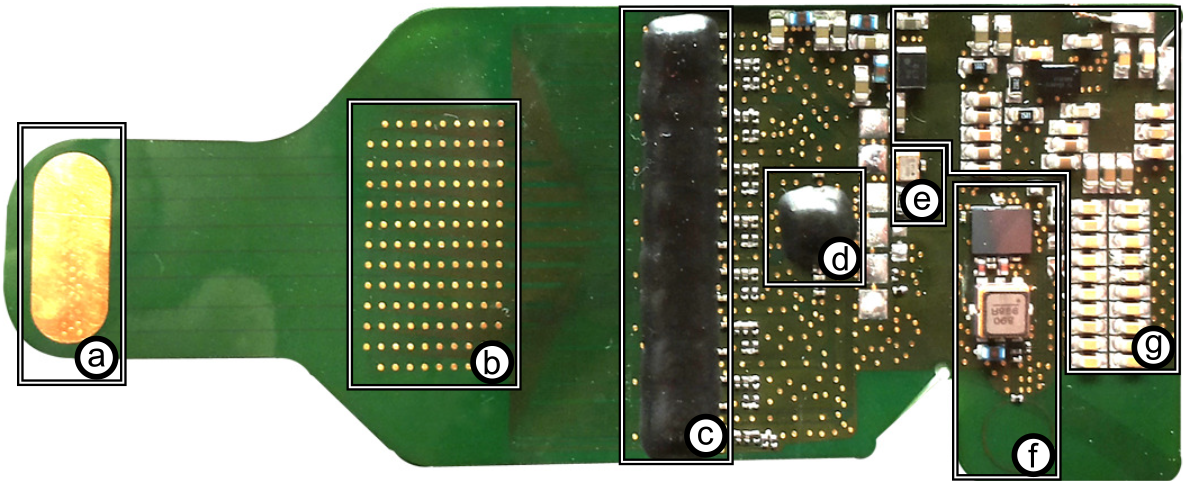
Implant prototype: (a) Reference electrode, (b) 128 electrodes, (c) 8×RHA, (d) ASIC, (e) 24 MHz clock, (f) RF-transceiver, (g) Inductive energy link

**Figure 6:**
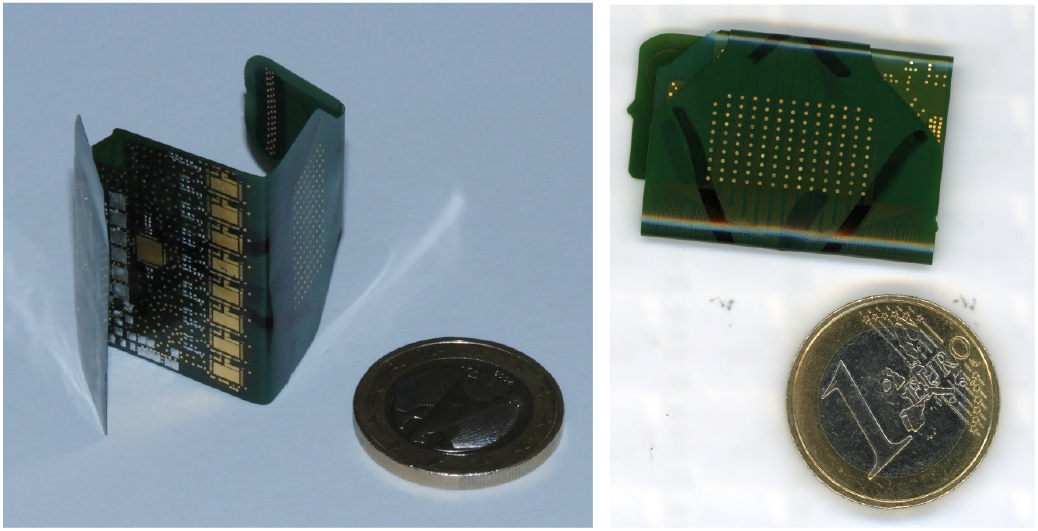
The implant is based on a flexible substrate, which allows it to be folded along three lines. This reduces the required space for implantation

**Figure 7:**
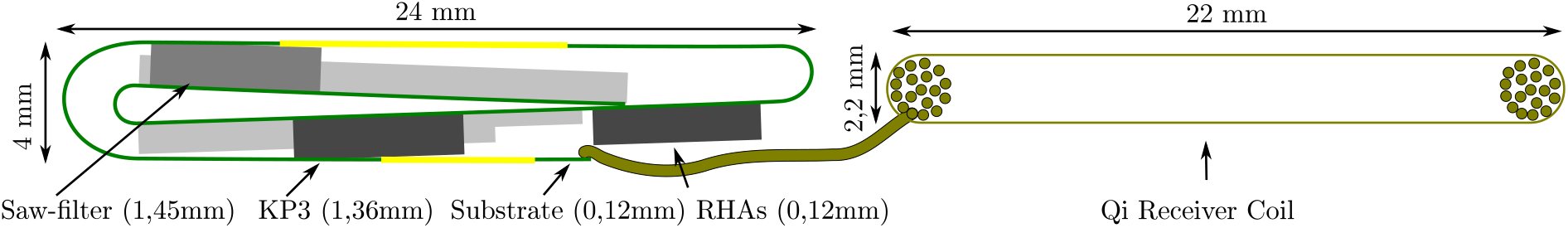
Kalomed Prototype with bending radius of 0.64mm (left) and coil for power supply (right)

**Analog front-end:** For the analog front-end Intan RHA2116 chips are used, which include the neural amplifiers and an analog-to-digital converter (ADC). Eight of these ICs are part of one implant, where each RHA provides 16 analog channels. The RHA contains, beside bio-signal amplifiers with a band-pass filter, an ADC that allows sampling all its individual channels at 10 kHz and 16 bit resolution. The integrated ADC is documented in a previous version of the RHA2116 data sheet. This part of the description was removed from the actual documentation. For newer designs, it is suggested to use the Intan RHD2132. We chose to operate the ADCs with their full 10 kHz sample rate which allows to reuse this setup with intracortical electrodes for recording action potentials. These ADCs generate 8 parallel data streams with a total of 20.48 MBit/s. On the other end of the data processing chain, the RF transceiver is only capable of transmitting up to 0.515 MBit /s.

**ASIC:** The 8 ADC data streams are collected by an in-house designed digital ASIC [42]. Besides serializing these parallel inputs, the ASIC has the capability to significantly reduce the incoming data according to user-defined parameters such as sample rate, resolution and the selection of electrodes which are included in the recording. Since the performance of the implanted electrodes can degrade with time, all the parameters can be changed dynamically during runtime in order to utilize the limited RF data bandwidth in an optimal way. Thus the implant uses the bi-directional nature of the RF link for receiving user-control commands from the base station during operation. The ASIC also controls the RF transceiver (e.g. initializing the connection and its parameters) as well as provides and caches the outgoing data (embedded into a suitable and compact transmission protocol) for achieving a high and continuous data transmission rate via the SPI connection. Furthermore, the ASIC contains integrated test-pattern generators, which can be used instead of the real measurement data from the Intan RHA2116 ICs.

**Nano FPGA:** Taking the data processing structures from the ASIC, we re-implemented the design in a way suitable for Microsemi IGLOO AGLN250 nano field programmable gate arrays. This allows us to provide a complete neuro-implant development system as open source solution exclusively using off-the-shelf components. Besides implementing the firmware for the Intan RHA analog-front end, we also wrote a second version for the newer Intan RHDs. Table 1 shows the required resources on the FPGA. The bare die of the FPGA is only slightly bigger (3.22mm × 3.48mm) compared with the ASIC but has a larger buffer for avoiding data loss during data transmission pauses. However, this component is still small enough to be used on a neuro-implant development system with the same size. In future designs of our implant development prototype, the nano FPGA will allow us to develop and test new versions of the data processing while keeping the test system in realistic dimensions. We provide the RHA and RHD based nano FPGA firmwares as well as tests boards for the Microsemi nano FPGA and the Intan RHD2132 as open source.

**Table 1:** Usage of the IGLOO nano FPGA resources for an implant with Intan RHA or RHD analog front-end. A large portion (up to 33% in the case with RHAs) of these core resources are by optional virtual RHAs/ RHDs for testing purposes.

| Resource | Usage (RHA) | Usage (RHD) |
| --- | --- | --- |
| CORE | 5859 of 6144 (95%) | 5236 of 6144 (85%) |
| IO (W/ clocks) | 38 of 68 (56%) | 38 of 68 (56%) |
| GLOBAL (Chip+Quadrant) | 6 of 18 (33%) | 6 of 18 (33%) |
| PLL | 0 of 1 (0%) | 0 of 1 (0%) |
| RAM/FIFO | 8 of 8 (100%) | 8 of 8 (100%) |

**Problems with flexible PCBs:** Due to problems with the quality of the PCB foils (shorts created by shifts between the layers of the foil during production), the implant prototype used for testing was equipped with only one instead of all 8 amplifier arrays. This has an effect on the power consumption (each RHA array consumes 5mW in this scenario) and consequently on the number of available channels for measurements. For testing different configurations, the 16 physically available measurement channels can be combined with the internal RHA test pattern generators in our ASIC. Besides the missing RHA arrays, the prototype supports all functions of the final implant, especially the complete wireless power and data transmission.

### Performance of the wireless module

Examples boards with the wireless module were produced on 150*μ*m thick FR4 and 50*μ*m thick flexible PCB-foil substrates. Both versions were tested successfully. However, due to the required very fine resolution (50 *μ*m strip width and distance between elements) of the PCBs, most of the flexible PCB-foils were produced with faults (e.g. shortcuts). Fortunately, we were able to fix some of them by manual cutting and grinding.

**Power link:** For the secondary side, we used a handwound coil (see figure 3) with 20 mm × 20 mm size and 18 turns of litz wire (20 × 0.05 mm individual wires). For the power-receiver IC we used, the maximum distance is defined by the Qi standard (version 1.0) with 5mm. This requires that the secondary coil is placed between skin and skull with two thin wires through the skull. Our transmitter can bridge a distance of 4.5mm with the described coil (L=10.5uH, Q=1). With a modified receiver coil we reached 5.5mm (L=15uH, Q=0.76). Figure 8 shows results of a range measurement, and how the base station adjusts the field strength and frequency depending on the distance and the inductance of the receiver coil. The figure shows no significant difference in the operation point (frequency and voltage) for air or meat as transmission medium. The received rectified power was set to 100mW for the experiment.

**Figure 8:**
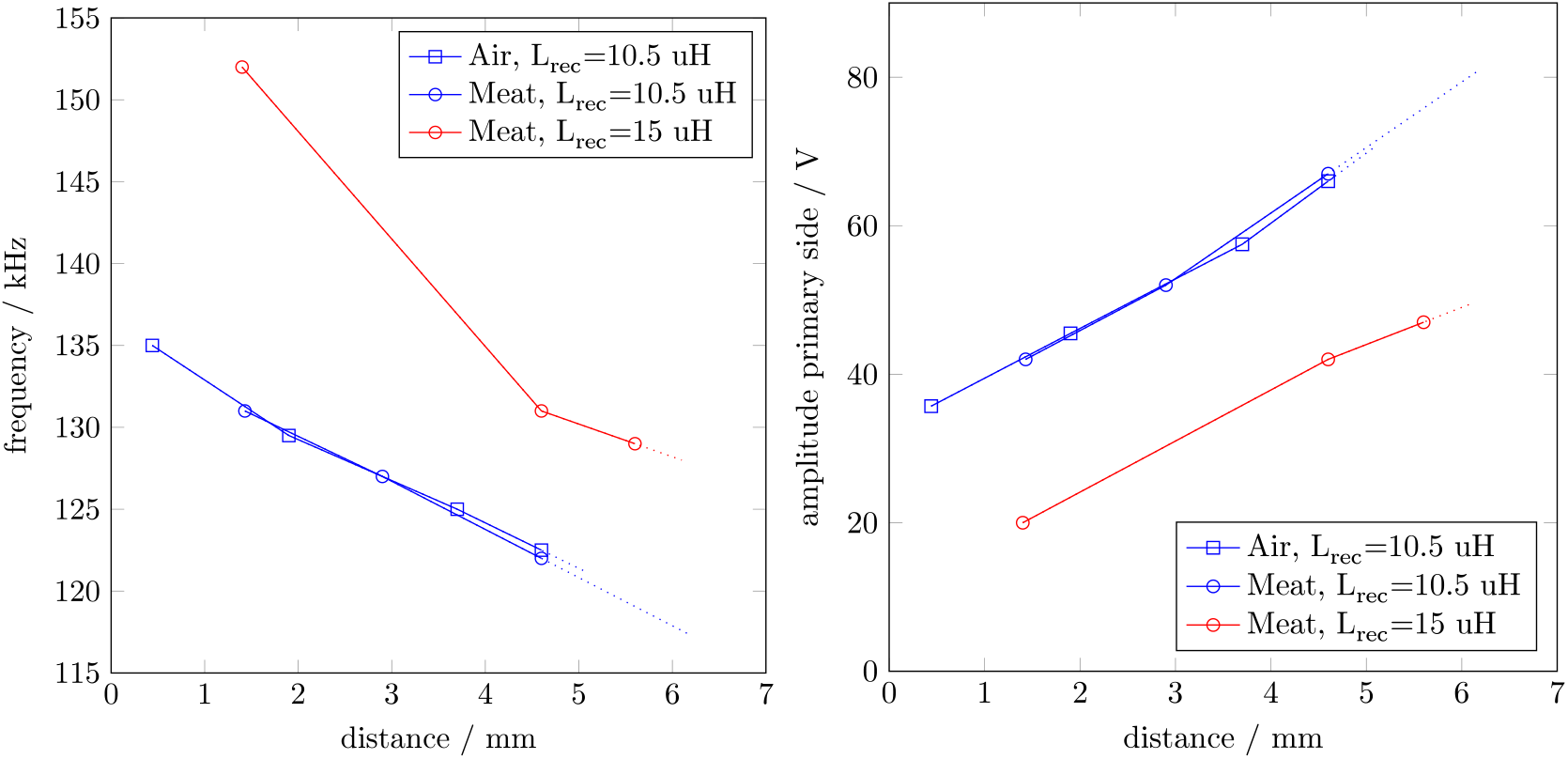
Wireless operation distances and according frequencies and primary voltages.

An update of the Qi standard [55] was announced, which will work over distances between 12mm and 45mm while being backwards compatible with the existing receivers. Furthermore, the 'Rezence' standard from the alliance for wireless power was also announced to have similar properties. These new standards are based on magnetic resonance which permit thick obstacles between the primary and secondary coil. It is expected that an update of our implant to these new standards will allow to place the secondary coil also under the skull.

**Data link:** We measured data transfer rates with the implant prototype in wireless operation (Figure 9). For the measurement we configured the implant to sample 52 channels at 2kHz and a resolution of 10 bits, which generates a data stream on the implant with 1.12 Mbit/s while the Microsemi transceiver shows a limitation of 515 kbit/s. This guarantees a full TX-buffer and allows to measure the maximal transmission performance. For each measurement condition (medium and distance), a set of 10 data tracks was recorded, each containing 100,000 sample sets. The duration of the transmission for each set was measured to reach the transmission rate.

**Figure 9:**
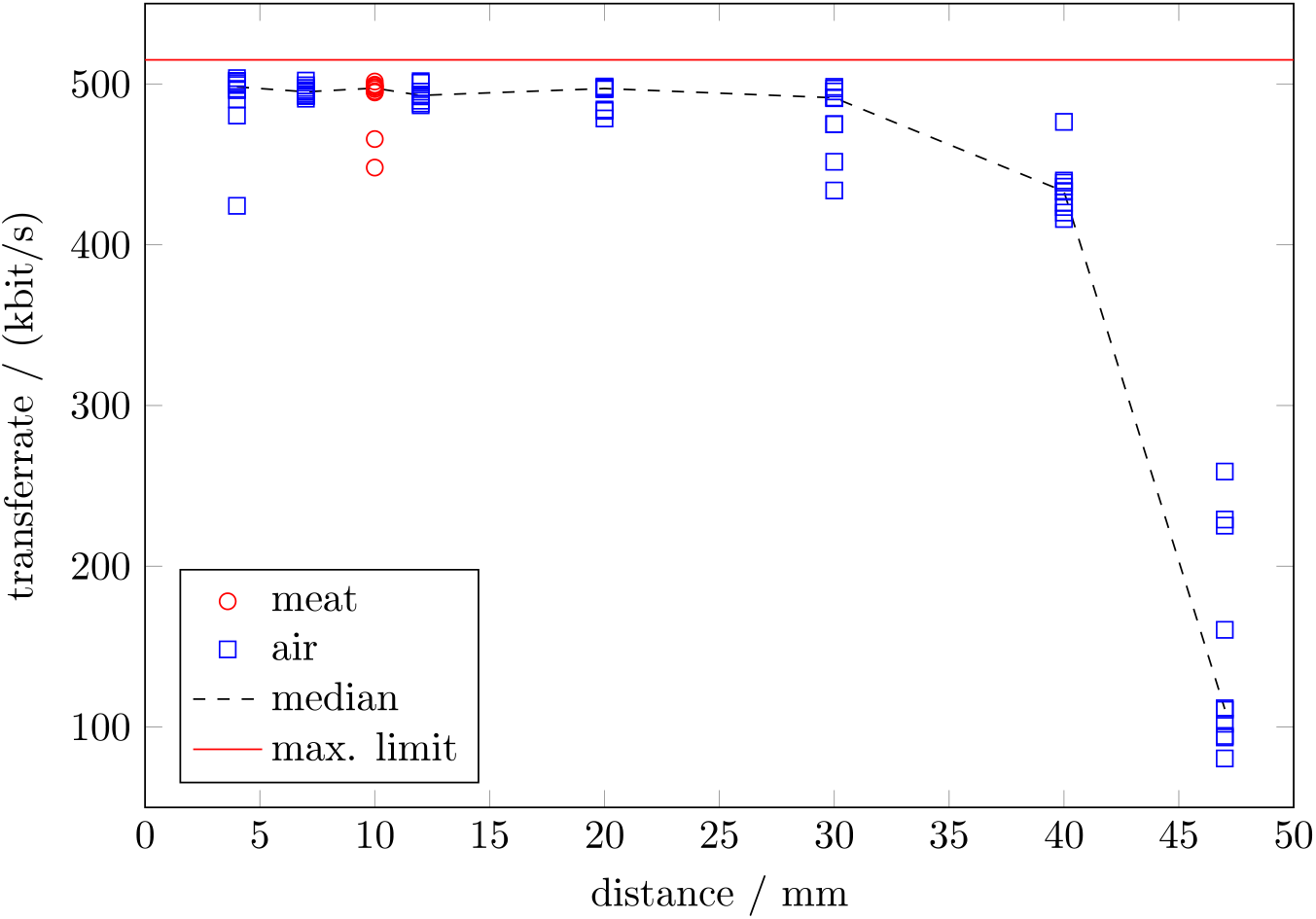
Data transfer rates for different distances.

For simulating in-vivo-measurements, we placed the implant prototype between two 1 cm thick stacks of sliced meat. We also tested the implant in air and observed comparable transmission rates at similar distances. Most important is the result that the data can be transmitted with almost maximum transceiver speed through 10mm of meat. A data transmission was possible up to 47 mm, but with a strongly reduced data rate due to the re-transmission of corrupted packets. Also under good conditions some samples are lost, because the Microsemi transceiver is not optimized for real time transfer but for good data integrity. The reason for the data loss lies in the limited amount of buffer the ZL70102 owns. In the case that this transceiver’s buffer is filled with a constant data rate, even small transmission pauses will fill the ZL70102 buffer completely in a short amount of time. Data needs to be discarded if it can not be buffered elsewhere on the implant. Time-stamps are included by our implant electronics to reconstruct the timing, even if packets are lost.

### Performance of the analog front end

For a rms-noise test, we analyzed the signals from a measurement, where the electrode array of the implant prototype was placed in Ringer solution (Figure 10). Since many of the externally produced PCB-foils had defects, we decided to use a repairable sample with only one Intan RHA2116 chip with 16 working channels. The prototype implant (with one RHA) was working in wireless operation, sampling 16 channels with 1kHz and a resolution of 10 bits. The rms noise of the measurement is 7.9*μ*V. Figure 11 shows the spectrum of the system noise. Figure 12 shows what sinusoidal waves look like when recorded with the analog front end of the implant.

**Figure 10:**
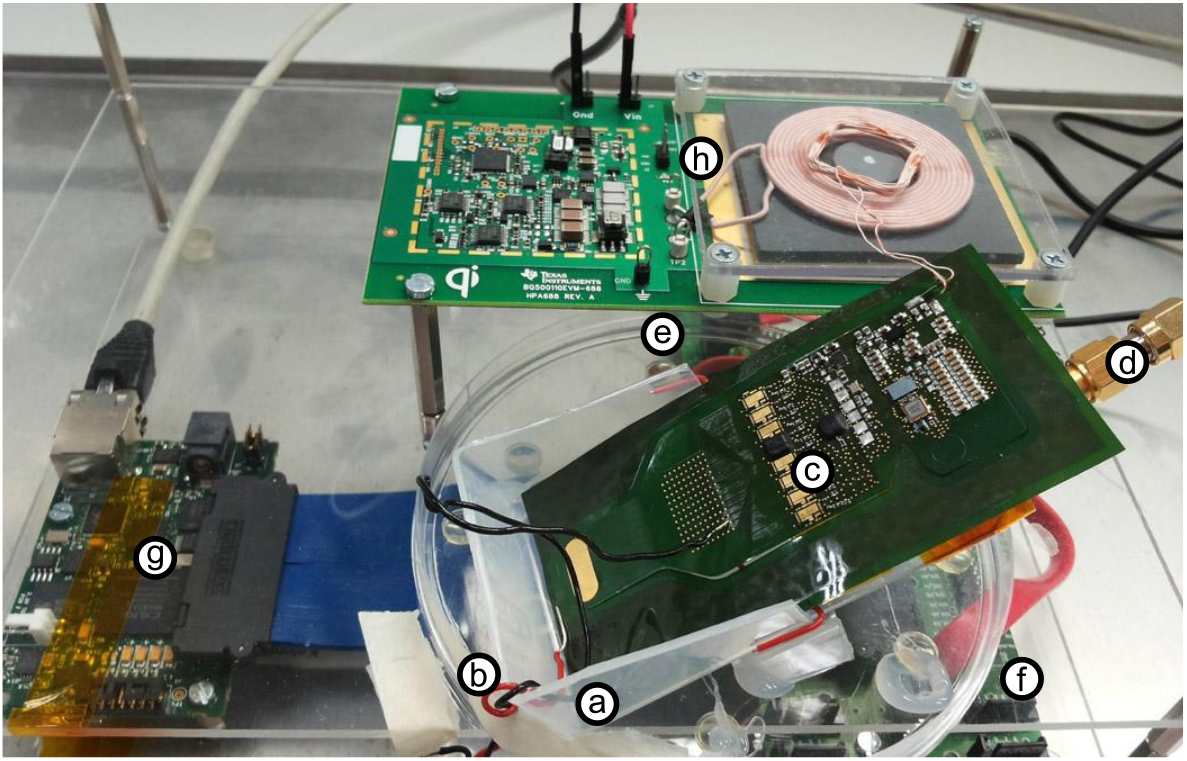
The complete system and measurement setup. A plastic box (a) is filled with Ringer Solution to simulate the fluids around the brain. The red and the black wire (b) are dipped into the fluid to apply a test stimulus between the electrodes and the reference electrode of the implant prototype (c). Underneath the implant lies the receiver antenna which is connected (d) to the base station receiver board (e). An adapter board (f) connects the receiver to the base station FPGA board (Orange Tree ZestET1) (g), which provides the data via Ethernet. The implant is powered using the TI bq25046EVM-687 kit board (e).

**Figure 11:**
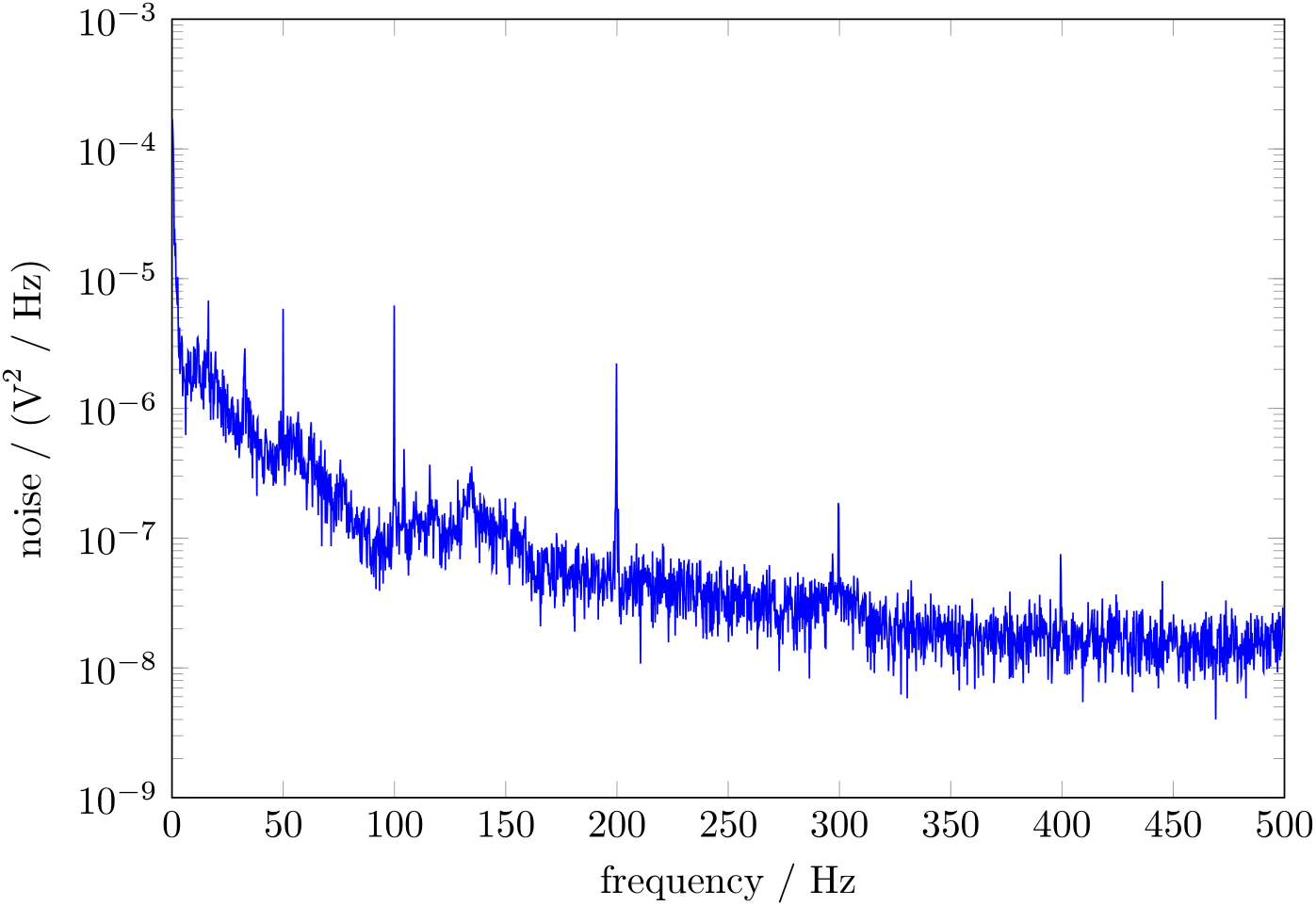
Noise spectrum for open inputs in Ringer solution.

**Figure 12:**
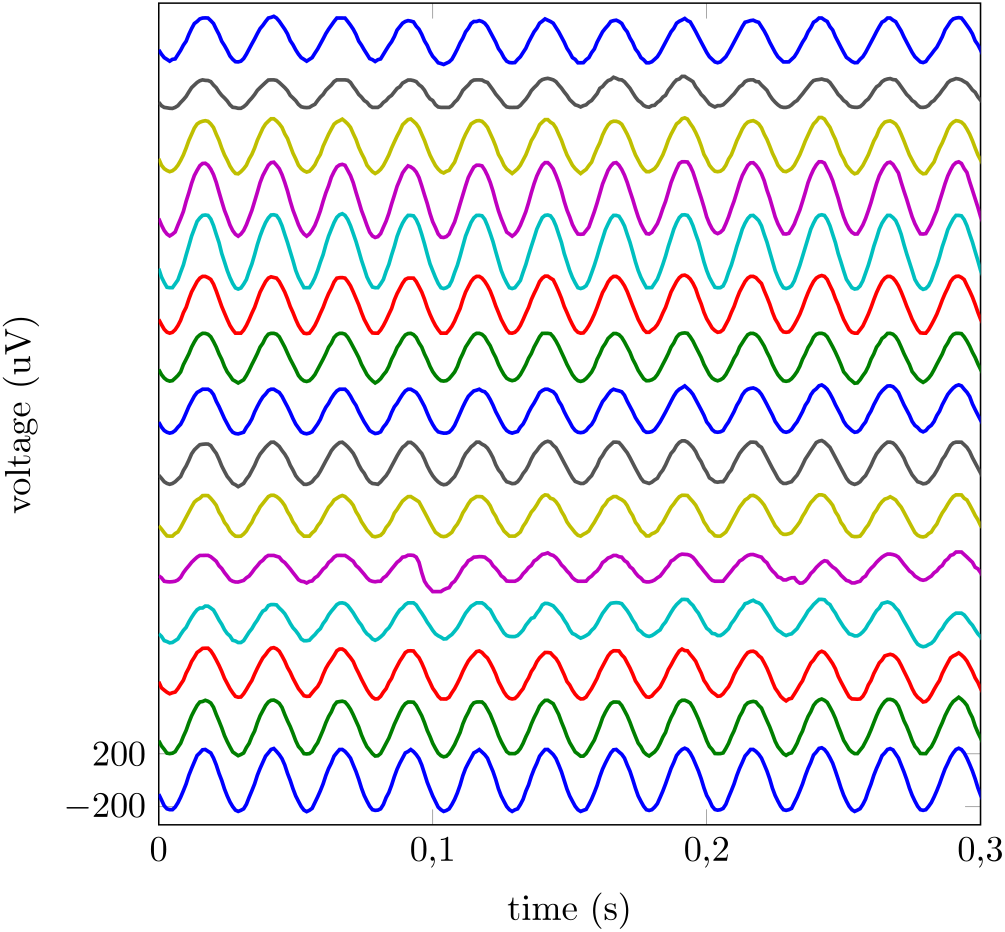
Received signals for a 40 Hz test signal. The amplitude depends on the distance between the stimulating wire and the channel electrode. The different channels are depicted in offset steps of 400 μV.

**Tests in a real-world application:** First successful tests of the electronics on non-miniaturized, non-wireless test boards were conducted in an animal experiment. Our goal was to test if the functionality can be demonstrated under real-world conditions. We restricted our experiments in line with the 3R-rules (reduce, refine, replace) to the recording sessions required for that purpose. In a first test we connected the system via cables to electrodes which were already implanted in an awake behaving animal (Macaca Mulatta) for a series of other neuro-research experiments. These implanted electrodes [44] were based on the substrate which we planned to use for the next generation of our flexible ECoG-implant.

We tried to repeat this with the miniaturized and fully wireless implant on the flexible substrate. We soldered cables onto the individual electrodes of the implant. Similar to the first test, we plugged the implant into the connector of a pre-implanted surface electrode grid. In contrast to the first test, a problem with the system was revealed. In the case that the reference electrode of wireless system was not grounded, the amplitude of the recorded signal was strongly reduced and the neuronal signal nearly vanishes from the recorded time series. The reason for this behavior is not fully understood yet.

One hypotheses is that this configuration allows the wireless power supply to induce an additional, strong 100kHz sinusoidal signal on top of the neuronal signal at the inputs of the amplifier. These combined signals are now larger than the threshold voltages of the Intan RHA’s ESD protection diodes of the analog input channels. As result, the protection diodes open a direct connection to electrical ground which eradicates the signal.

In tests without animals, measurements with an oscilloscope of voltages differences between the RHA’s analog inputs and its reference revealed such voltages. A larger distance between the RHAs and the energy transmitter/ energy harvesting coil may reduce the problem or switching to a different kind of wireless energy link system might also remove this problem. We were able to confirm that the wireless power supply still functions if the distance between the receiver and the coil is increased to even 10cm. It has to be noted, that this problem could also be an artifact of the several tenth of cm long cables soldered onto the electrodes for allowing the implant to be connected to the already implanted electrode-grid. These cables could possibly act as an antennas which allow the induction of these voltages. However, before we could determine reason of the problem or find a cure, the financial support for the project ended.

### Estimated power consumption of the implant

An estimate for the main electrical loads of the components of the implant are shown in table 2. Combined with the losses of the power supply ICs, the fully equipped implant will dissipate about 110mW-140mW of power, while the intensively tested prototype with one RHA consumes about 73mW-103mW (both are presented in the next section). For safety reasons, the power receiver IC is programmed not to accept more than 200mW to limit the production of heat in a case of failure.

**Table 2:** Estimated power consumption of the implant’s components.

| Component | power consumption |
| --- | --- |
| Microsemi ZL70102 transceiver | 17mW (measured) |
| ASIC | up to 9.44mW (measured) |
| Clock quartz | 16.5mW (measured) |
| RHA amplifier arrays | 5mW (each IC) (measured) |
| DC/DC-Converter | 8.5mW (for 1 RHA), 15mW (for 8 RHAs), (data-sheet) |
| TI inductive power receiver | 10-40 mW (data-sheet) |

### Examining the implant’s thermal properties

**Tissue Heating:** A major concern for neural implants is the heating of the tissue, as proteins start denaturation at approximately 40°C. The IEEE Standard [56] states a brain temperature 40.5°C as critical for a heat stroke. The tissue temperature close to the implant is affected by different heat sources. Most critical is the joule heating of the implant electronics due to the high power densities (e.g. 17mW in 9*mm*^3^ for the transceiver IC). Due to the folded structure of our implant, all active components are embedded inside the implant, which strongly increases the contact area to the tissue.

Another heat source are eddy currents from the inductive field of the power and data transmission in the conductive tissue and in the implant. The eddy currents are expected to be negligibly small, according to the stable operating point shown in Figure 8. Heating by the field of the data transmission in the MICS-band can also be neglected. The whole RF transceiver only consumes 17mW of power, only a percentage of it is really transformed into field energy.

Finally the joule heating of the base station coil, which has contact to the skin above the implant, increases the temperature of the tissue. A simple countermeasure could be a cooling system over the implant attached on the outside of the body.

**Simulation of joule heating:** As our prototype is equipped with only one amplifier array instead of eight, we used a simple FEM model (COMSOL) to evaluate the heating of the final, fully assembled and folded implant. We used the outer dimensions shown in Figure 7 and applied a heat source of 100 mW distributed over the volume of the implant. We chose 100mW for the simulation as an estimation for the typical power dissipated by a fully equipped implant in operation. Actual values might be lower or higher depending on the number of active RHAs and the actual efficiency of the TI inductive power receiver, but they do not change the magnitude of the resulting temperatures. Figure 13 shows the heat-up curves at the surface of the implant and at different distances within living tissue. The temperature at the surface in thermal equilibrium is calculated to be 0.25K above the starting temperature of 37°C, in a sphere with 10cm diameter and a border temperature of 37°C. Additionally, Figure 14 shows a temperature map taken after 300 seconds, with the 24mm × 4mm cross section of the implant positioned in the center.

**Figure 13:**
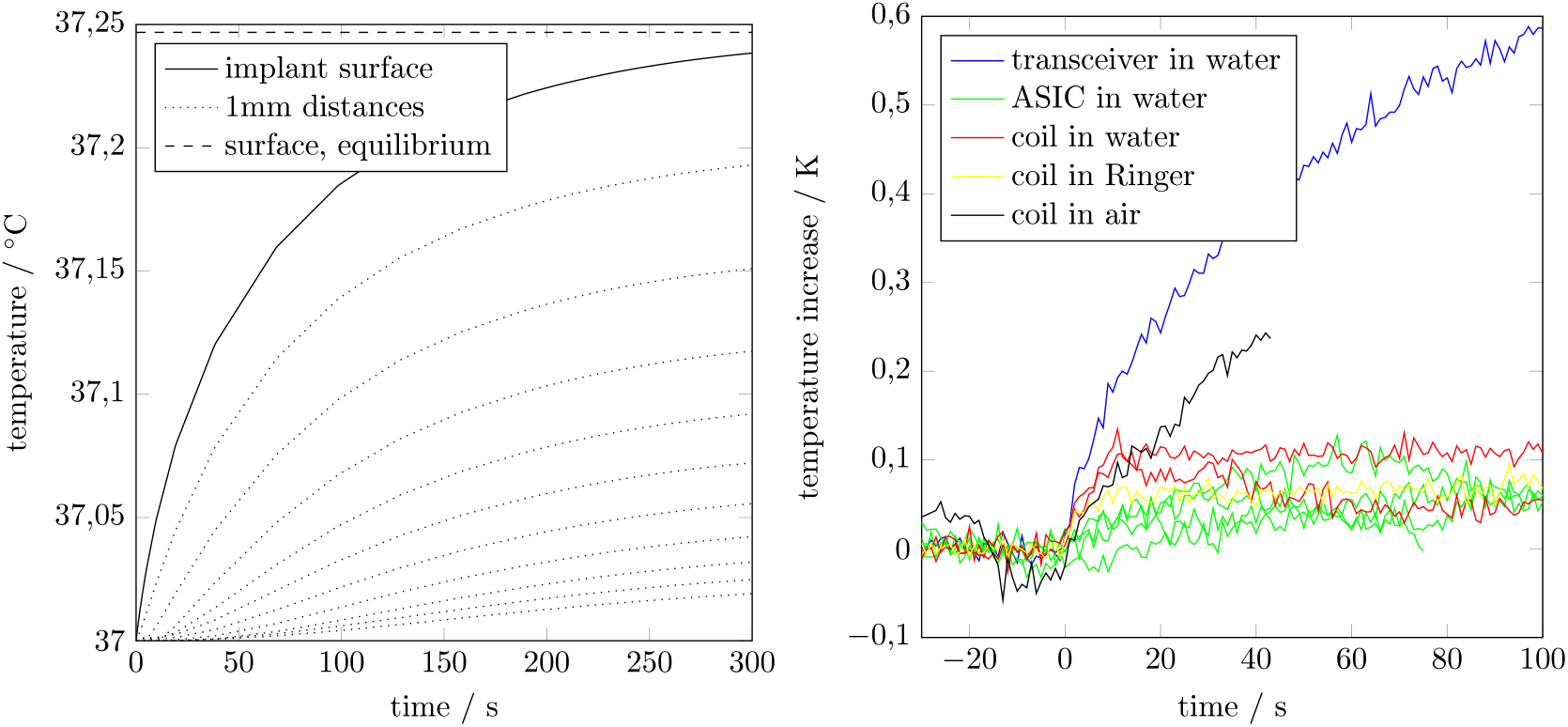
Left: Simulated heat-up. Right: Measured temperature increase after power on.

**Figure 14:**
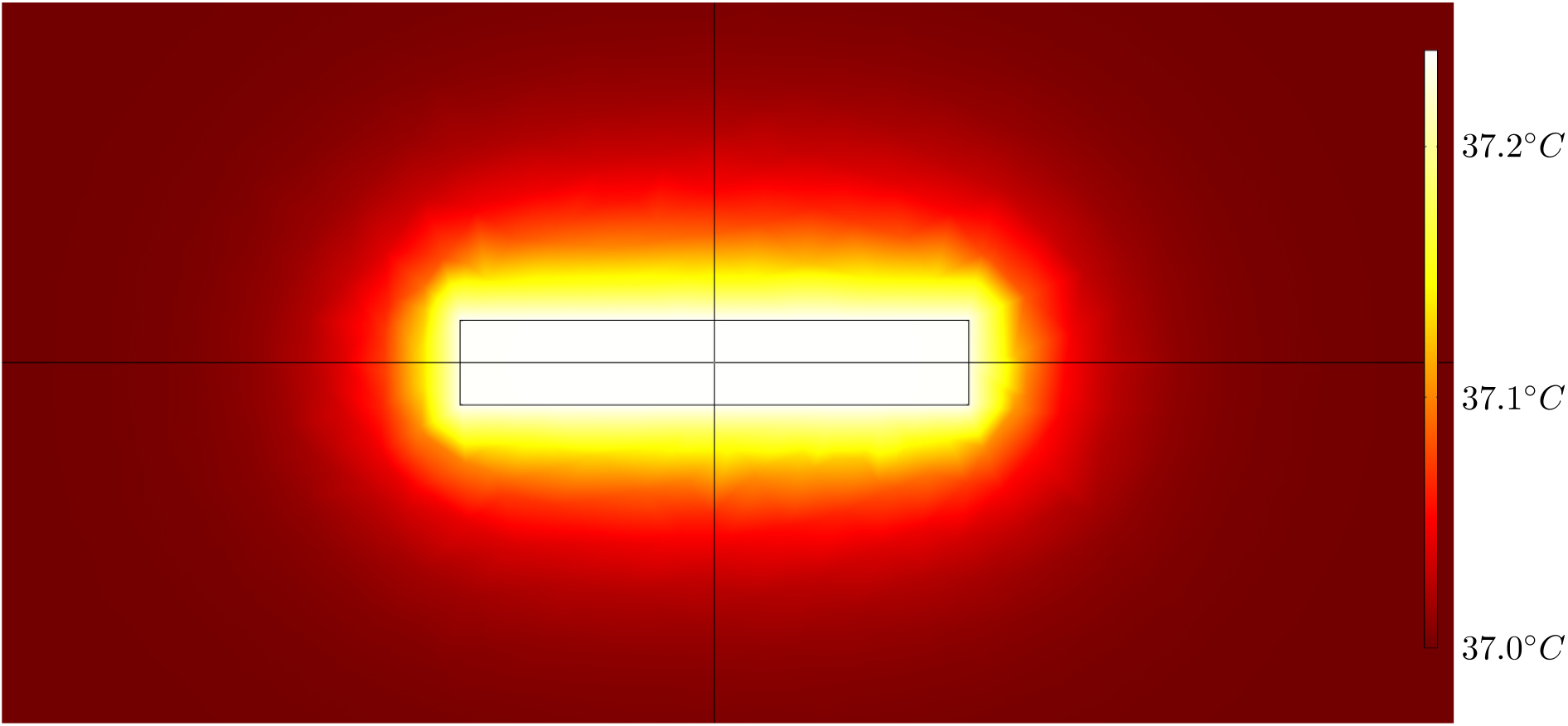
Temperature distribution after 300 seconds, calculated with a simple FEM model (COMSOL). Rectangle shows the 24mm × 4mm implant cross section dimensions.

**Measurement of total heating for the prototype:** In addition to the simulation, we made an experiment to observe the heating in wireless operation. The measurement setup is shown in Figure 15. The implant prototype was isolated with a thin PCB-foil of plastic wrap against a liquid medium (Ringer solution) with a volume of 150ml. For the temperature measurement we used a thermocouple and contacted it to different parts of the implant. Based on the simulation, we expected the surface temperature to be saturated after a few minutes. Longer test periods are not expected to provide meaningful results, because in contrast to a living subject, the surrounding tissue (in our case 150ml fluid) would heat up more and more because of the small volume and lack cooling by blood perfusion.

**Figure 15:**
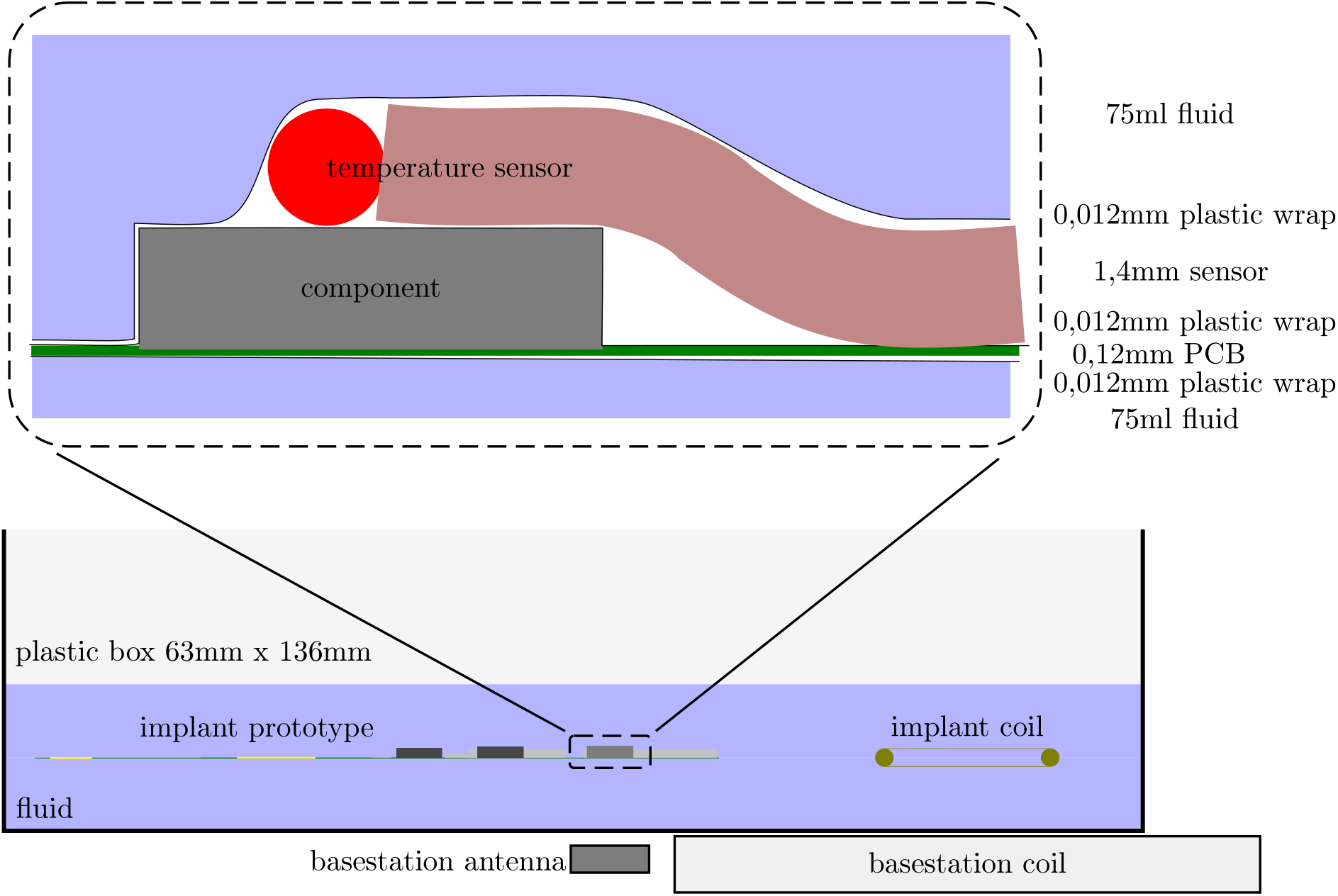
Measurement setup for testing the heating up of the implant.

The black curve in Figure 13 shows a rapid joule heating of the coil in air, while the heating saturates at approximately 0.1°C in water (red) and at even lower values in the conductive Ringer solution (yellow). Close to the ASIC, which is covered under a 0.75*mm* plastic housing and has a power dissipation of 9.44*mW*, we also measured a temperature increase below 0.1°C.

In contrast to the low heating at the ASIC, the blue curve shows the temperature on top of the unencapsulated RF transceiver IC, which has a power dissipation of 17 mW. In Figure 7, the transceiver is located behind the saw filter, which is ca. 1mm higher than the transceiver IC. Thus, after folding, the transceiver IC has no direct contact to the tissue. In our experiment, the PCB-foil was not folded and we saw a strong temperature increase at the contact between the IC and the liquid medium. The ground planes in the substrate are expected to distribute the thermal power more equally to the outer implant surface.

## Discussion

This article starts with the question if it is possible to design a fully wireless neuroimplant and it’s external base-station such that the results can be reused by other groups as starting point for their own technology development activities. As an answer, we present a system concept which can be transferred into real hardware by using only commercial off-the-shelf components. This hardware realization is explained in detail and its performance reported. Furthermore, an analysis concerning the heat development of the implant was conducted in measurements and simulations. Finally, all design files (circuit diagrams, boards, firmwares, and software as well as documentation concerning the development process) are made open source in the supplemental materials. We deliver two versions of firmwares for the Microsemi IGLOO nano FPGA. One was written for the Intan RHA2116, using undocumented ADC functionality, like the ASIC we used for the measurements with the flexible implant prototype and other firmware version which was optimized for using the newer Intan RHD2132.

**Comparison with other neuro-implants:** As a very important key-technology for future neuro-prostheses systems, building implantable systems is getting more and more popular. In contrast to other ‘typical’ systems, with our 128 channel system for measuring ECoG signals we developed an implant that can be placed completely under the skull and avoids energy storage elements (e.g. batteries) with a limited lifetime. We are convinced that it is very high important for long-term stability and safety that the skull can be closed completely again after implantation. This keeps the natural barrier against germs intact and prevents cerebral fluid leakages [28].

Another approach taken by [57] is to replace parts of the skull directly with the implant or components of the external base-station [58] but this leaves the skull constantly open.

A completely different approach was chosen by [59, 35]. They developed a 100 channel recording unit (LFPs and action potentials) which can be implanted into the torso like a pacemaker. The system uses RF (3.2/3.8 GHz) and infra-red light for the wireless data transmission. The power is provided by a Li-ion battery and can be recharged with 2MHz electromagnetic waves. A similar approach is presented in [60]. There they use Bluetooth for exchanging data, a battery, and titan casing for recording from two times 64 channels. Such a battery has a limited lifetime which is clearly below the required several decades. This strategy can’t be transferred to implants that will be installed under the skull. One major problem of neuro-implants, in comparisons e.g. to pacemakers, is that after some time scar-tissue or even bone encapsulates the part of the implant that is situated within the head. Without damaging the brain tissue, replacing the implant or parts of it (e.g. batteries) becomes very problematic. [61] presents a two times 32 channel recording system. This system is designed such that each of the 32 channels are implanted into separate cortical areas. It has inductive power supply and infra-red based data transmission. Part of the system is implanted under the skull and connected with wires to a data processing unit which is installed under the skin. [62] is an earlier version of [61], which again seems to be the predecessor of [35].

As equipment for research applications, we find a recording system for 32 channels using frequency-shift keying (FSK) modulation in the 4GHz range for data transmission and is powered by a battery [63]. This system is large (38mm × 38mm × 51mm in an aluminum enclosure) and installed outside of the subject. Also a lot of research effort went into the development of recording systems which can be put e.g. on top of freely moving or even flying insects. [64] demonstrates a four channel system that uses a 900MHz RF data link and a battery for providing power to the system. The weight of the system is very low. [65] presents a similar system which is able to record 10 channels with 26.1K samples per second and 4 additional channels with 1.63k samples per second. For this system a battery is not required because it harvests energy from RF electromagnetic waves. [66] shows a device that can apply electrical stimulation to the central nervous system of a large beetle and control it.

The development of a fully implantable system based on the Utah needle array is shown in [67, 68, 69, 70]. This system contains individual threshold-based action potential detectors for all its 100 channels. It sends the detected spikes via a 900MHz data link to its base station. However, the system was tailored for recording the timing of action potentials and not of ECoG signals. For only one selected channel, the device is able to deliver the recorded time series with 15.7k samples per second. Energy is also provided via an inductive link. Also it is not yet clear if the approach with these needle arrays will work over several decades [71]. Our own experience shows that recording with surface electrode mats are much more long-term stable than with needle arrays.

In [72] the development of an integrated circuit for recording 128-channels with on-the-fly spike feature extraction and wireless telemetry is presented. It uses for data transmission UWB (ultra wide band with 90Mbit per second) and has no solution concerning the power supply. [73] presents an all-in-one chip solution for 32 recording channels. It is able to collect power via a 13.56MHz inductive link. Data is transmitted via 900MHz FSK coded. In [74] a complete chip-set for a 100 channel recording system is shown. It has an wireless data transfer unit and harvests energy via an inductive power link.

Concerning the individual components of such an implant [75, 76, 77] like e.g. bio-signal amplifiers [78, 79, 80], analog-to-digital converter [81, 82, 83], data processors [42, 84], wireless data transfer sub-system [85, 86, 87, 88, 89, 90, 91, 92, 61], or energy harvesting [93, 94] a large number of publications exist.

Many of those components, system parts, and system designs show remarkable performances but they are not freely available. The implant we present has lower specs compared to these highly optimized solutions but our system can be re-build by everybody and then be modified to your heart’s content.

**Bio-compatibility:** One of the remaining obstacle preventing us from applying the system in-vivo, are long-term stable bio-compatible coatings which can protect the electronics from the harsh fluidic environment in the body. This coating has to stay intact over many years and has to be very thin and flexible. We designed the implant to be completely coated in a first processing step. In a second step, the coating must be removed from the electrodes and then the implant is folded at three folding lines for reducing the required area. Therefore, it is required that the coating is not only flexible but has a good adhesion to all the components. We expect that the adhesion between the coating (e.g. Parylene C) and the used material for the PCB-foil (DuPont Pyralux AP and insulating resist) might cause problems. This requires to change the substrate of the PCB-foil to something more suitable (e.g. Parylene C as well) and will hopefully give us the opportunity to reduce the thickness of the substrate for improving the bending radius of the PCB-foil. Currently, we are testing several promising candidates for coatings and substrates [43].

**Electrical stimulation:** Another goal for the future is to add comprehensive stimulation capabilities to the implant for allowing electrical stimulation of the brain tissue (e.g. for visual cortical prostheses). Our ASIC has the capability of stimulation. However, it can only create simple 3.3V uni-polar stimulation pulses (unregulated in current strength) on 8 extra pads. This was our very first step of including stimulation into a design. For the implant on the flexible substrate, we decided not to connect these outputs to the electrodes. Instead we focused our efforts [95] on developing better current pumps optimized for the high-voltage electrical stimulation with up to ±90V and ±10mA [96], which is required for ultra high density surface grids with very small electrodes. In the case of stimulation with such high voltages, it is important to protect the sensitive analog-inputs of the recording system. High voltages can destroy the bio-signal amplifier which have a working range of several around 100mV. Even in the best case scenario, these amplifiers are overloaded and this will cause severe recording artifacts for many 10ms. In addition, the ESD protection diodes of the amplifiers’ input channels will kick in and burn an undefined amount of current from the stimulation pulse. A solution against this problem lies in fast analog switches which can withstand such high voltages while keeping the distortion of the sub-mV neuronal signals as low as possible. We are also working on this kind of optimized switches for this special application [97].

**Possible improvements for the design:** Especially two aspects of the implant need improvement in future: 1.) By exchanging the power harvesting to magnetic resonance technology (the new Qi standard or the Rezence wireless power charging standard), the maximal operating distance between the primary and secondary coil can be increase to up to 40mm. In the actual state, our implant requires an energy harvesting coil between the skin and outside of the skull which is connected with two small wires to the implant under the skull. 2.) The effective data transmission rate is limited to 515kbit/s. For many applications this transmission rate is too low. We looked into the possibility of optical data transmission by infra-red light. Together with the BIAS (Bremen institute of applied beam technology), we tested the feasibility of this idea by sending high-frequency signals through meat, skin and bones. We expect that data transfer rates of over 100MBit/s should be possible with optimized micro-optics and a vertical-cavity surface-emitting laser (VCSEL) on the implant as well as a suitable external receiver. If code division multiple access (CDMA) is used, even several implants can send information on the same wavelength. For the channel from the external base-station to the implant, the slow RF connection still can be used or also replaced by an IR data transmission (which is more challenging) on a different wavelength.

Ex-vivo tests have proven the feasibility of our system design. However, during first *in vivo* functionality tests, we ran into problems (for details see results) which never were seen during lab-bench tests. As a result, the amplitude of the measured neuronal signals nearly vanishes when the reference of the Intan RHA2116 and the base-station doesn’t have a low impedance connection. Such a cable is not a requirement we want for a wireless system. The reason for this is still unclear and maybe an artifact of the very special measurement setup (e.g. long cables soldered onto the electrodes of the flexible implant as a connection to an additional electrode which was already implanted in the animal) or the close distance between the energy coil placement and the rest of the implant. After solving this problem, real *in vivo* tests need to be performed in order to verify the system performance under real measurement conditions. This information is required to estimate the development steps that have to be taken for making the system safe enough for using it for human patients.

**In summary**, the actual state of the implant is not yet ready for implantation, especially not for long-term implantation in medical applications. Several problems have still to be solved in future development. Nevertheless, we deliver an open source tool kit completely based on commercial off-the-shelf components. This collection contains circuit diagrams, board designs, FPGA firmwares, and software which allows interested researchers to develop their own wireless neuro-implant without starting from scratch.

## Materials and Methods

All experimental procedures using animals were approved by the local authorities (Der Senator fuer Gesundheit, November 11 2014) and were in accordance with the 3R-priciples, the regulation for the welfare of experimental animals, issued by the federal government of Germany and with the guidelines of the European Union (Council Directive 2010/63/EU) for care and use of laboratory animals.

## Supplementary

In the supplemental data we present the design files for the firmwares, software and PCB designs as open source as well as documentation concerning the development process.

## Acknowledgements

We thank Norbert Hauser, Alexander Svojanovsky, and Mario Kaiser from Brain Products as well as Guido Widman and Christian Elger from the department of epileptology at the university hospital of Bonn for fruitful discussions. We thank Sunita Mandon and Tobias Tessmann from the University of Bremen for their support. This work was supported in part by Bundesministerium fuer Bildung und Forschung, Grant 01 EZ 0867 (Innovationswettbewerb Medizintechnik) and Grant 01 GQ 1106 (Bernstein Award Udo Ernst) as well as Research-Focus Neurotechnology University of Bremen, and the Creative Unit I-See ’The artificial eye: Chronic wireless interface to the visual cortex’ at the University of Bremen. Also this work was supported by the Deutsche Forschungsgemeinschaft priority program SPP 1665 ’Resolving and manipulating neuronal networks in the mammalian brain - from correlative to causal analysis’ (LA 1471/11-1).

### Conflict of interests

The authors declare that there is no conflict of interest regarding the publication of this paper.

### Author Contributions

D. R. and J.P. wrote the paper. K.R.P. and D.R. intiated and supervised the research in this project. D.R., J.H., J.P., W.L., D.P.D., S.P., K.R.P. and A.K. developed the system concept. J.P. and J.H. prepared and conducted the test and startup. J.P. performed the measurements. H.S. and A.K. performed the animal experiments. J.P. and J.H. developed the ASIC. D.R. designed the PCBs and PCB-foil for the implant, the wireless module as well as the base station. D.R. wrote the firmwares for the basestation’s FGPA and the nano FPGAs as well as the corresponding software package. D.B. worked on the wireless power transfer. S.P. provided the infrastructure for development, design and testing of the mixed signal circuitry. D.P.D. contributed the electronic design methodologies of the mixed signal circuitry. T.S. and M.S. developed the antennas. T.S., D.R., and M.S. created the corresponding antenna matching circuits. T.S. developed and built the energy harvesting coils. W.L. was responsible for the clean room technology. W.L. and E. T. contributed to the layout and realization of electrodes. D.G. contributed to the tests of the base station.

## References

[1] Peter H Schiller and Edward J Tehovnik. Visual prosthesis. Perception, 37:1529–1559, 2008.

[2] W. H. Dobelle. Artificial vision for the blind by connecting a television camera to the visual cortex. ASAIO Journal (American Society for Artificial Internal Organs), 46.1:3–9, 1999.

[3] E M Schmidt, M J Bak, F T Hambrecht, C V Kufta, DK O’Rourke, and P Valiabhanath. Feasibility of a visual prosthesis for the blind based on intracortical microstimulation of the visual cortex. Brain, 119:507–522, 1996.

[4] M. van Gerven, J. Farquhar, R. Schaefer, R. Vlek, J. Geuze, A. Nijholt, N. Ramsey, P. Haselager, L. Vuurpijl, S. Gielen, and P. Desain. The brain-computer interface cycle. Journal of Neural Engineering, 6, 2009.

[5] R.A. Andersen, E.J. Hwang, and G.H. Mulliken. Cognitive neural prosthetics. Annu. Rev. Psychol., 61:169–190, 2010.

[6] M.A. Lebedev and M.A. Nicolelis. Brain-machine interfaces: past, present and future. Trends Neurosci., 29(9):536–546, 2006.

[7] R.J. Ifft, M.A. Lebedev, and M.A. Nicolelis. Reprogramming movements: extraction of motor intentions from cortical ensemble activity when movement goals change. Front Neuroeng., 5(16), 2012.

[8] Wei Wang, Jennifer L. Collinger, Alan D. Degenhart, Elizabeth C. Tyler-Kabara, Andrew B. Schwartz, Daniel W. Moran, Douglas J. Weber, Brian Wodlinger, Ramana K. Vinjamuri, Robin C. Ashmore, John W. Kelly, and Michael L. Boninger. An electrocorticographic brain interface in an individual with tetraplegia. PloS one, 8(2):e55344, 2013.

[9] L.R. Hochberg, M.D. Serruya, G.M. Friehs, J.A. Mukand, M. Saleh, A.H. Caplan, A. Branner, D. Chen, R.D. Penn, and J.P. Donoghue. Neuronal ensemble control of prosthetic devices by a human with tetraplegia. Nature, 442:164–171, 2006.

[10] J.D. Simeral, S.-P. Kim, M.J. Black, J.P. Donoghue, and L. R. Hochberg. Neural control of cursor trajectory and click by a human with tetraplegia 1000 days after implant of an intracortical microelectrode. Journal of Neural Engineering, 2011.

[11] M.A. Lebedev, A.J. Tate, T.L. Hanson, Z. Li Z, J.E. O’Doherty, J.A. Winans, P.J. Ifft, K.Z. Zhuang, N.A. Fitzsimmons, D.A. Schwarz, A.M. Fuller, J.H. An, and M.A. Nicolelis. Future developments in brain-machine interface research. Clinics (Sao Paulo), 66(1):25–32, 2011.

[12] M. Velliste, S. Perel, M.C. Spalding, A.S. Whitford, and A.B. Schwartz. Cortical control of a prosthetic arm for self-feeding. Nature, 453:1098–1101, 2008.

[13] S. Musallam, B.D. Corneil, B. Greger, H. Scherberger, and R.A. Andersen. Cognitive control signals for neural prosthetics. Science, 305:258–262, 2004.

[14] D. Moran. Evolution of brain-computer interface: action potentials, local field potentials and electrocorticograms. Current Opinion in Neurobiology, 20(6):741–745, 2010.

[15] Beata Jarosiewicz, Anish A Sarma, Daniel Bacher, Nicolas Y Masse, John D Simeral, Brittany Sorice, Erin M Oakley, Christine Blabe, Chethan Pandarinath, Vikash Gilja, et al. Virtual typing by people with tetraplegia using a self-calibrating intracortical brain-computer interface. Science translational medicine, 7(313):313ra179–313ra179, 2015.

[16] David Rotermund, Udo A. Ernst, Sunita Mandon, Katja Taylor, Yulia Smiyukha, Andreas K. Kreiter, and Klaus R. Pawelzik. Toward high performance, weakly invasive brain computer interfaces using selective visual attention. The Journal of Neuroscience, 33(14):6001–6011, 2013 2013.

[17] Gerwin Schalk. Can electrocorticography (ecog) support robust and powerful brain-computer interfaces? Front Neuroengineering., 3:9, 2010.

[18] Tobias Pistohl, Andreas Schulze-Bonhage, Ad Aertsen, Carsten Mehring, and Tonio Ball. Decoding natural grasp types from human ecog. NeuroImage, 59:248–260, 2012.

[19] Tobias Pistohl, Tonio Ball, Andreas Schulze-Bonhage, Ad Aertsen, and Carsten Mehring. Prediction of arm movement trajectories from ecog-recordings in humans. Journal of Neuroscience Methods, 167:105–114, 2008.

[20] P. Brunner, A.L. Ritaccio, J.F. Emrich, H. Bischof, and G. Schalk. Rapid communication with a p300 matrix speller using electrocorticographic signals (ecog). Front. Neurosci, 5:5:doi: 10.3389/fnins.2011.00005, 2011.

[21] E.C. Leuthardt, G. Schalk, J.R. Wolpaw, J.G. Ojemann, and D.W. Moran. A brain-computer interface using electrocorticographic signals in humans. Journal of Neural Engineering, 1:63–71, 2004.

[22] D. Rotermund, U. Ernst, and K. Pawelzik. Towards on-line adaptation of neuro-prostheses with neuronal evaluation signals. Biological Cybernetics, 95:243–257, 2006.

[23] W. Wu and N.G. Hatsopoulos. Real-time decoding of nonstationary neural activity in motor cortex. IEEE Trans Neural Syst Rehabil Eng., 16(3):213–222, 2008.

[24] Z. Li, J.E. O’Doherty, M.A. Lebedev, and M.A. Nicolelis. Adaptive decoding for brain-machine interfaces through bayesian parameter updates. Neural Comput., 23(12):3162–3204, 2011.

[25] Kanber Mithat Silay, Catherine Dehollain, and Michel Declercq. Numerical analysis of temperature elevation in the head due to power dissipation in a cortical implant. Engineering in Medicine and Biology Society, 30th Annual International Conference of the IEEE, 2008.

[26] Sohee Kim, Prashant Tathireddy, Richard A. Normann, and Florian Solzbacher. In vitro and in vivo study of temperature increases in the brain due to a neural implant.” neural engineering. CNE’07, 2007.

[27] Sohee Kim, Prashant Tathireddy, Richard A. Normann, and Florian Solzbacher. Thermal impact of an active 3-d microelectrode array implanted in the brain. IEEE TRANSACTIONS ON NEURAL SYSTEMS AND REHABILITATION ENGINEERING, 15(4):493–501, 2007.

[28] J Voges, Y Waerzeggers, M Maarouf, R Lehrke, A Koulousakis, D Lenartz, and V Sturm. Deep-brain stimulation: long-term analysis of complications caused by hardware and surgery - experiences from a single centre. J Neurol Neurosung Psychiatry, 77(7):868–872, 2006.

[29] W. S. Lee, J. K. Lee, S. A. Lee, J. K. Kang, and T. S. Ko. Complications and results of subdural grid electrode implantation in epilepsy surgery. Surg Neurol, 54:346–351, 2000.

[30] Dileep R. Nair, Richard Burgess, Cameron C. McIntyre, and Hans Lueders. Chronic subdural electrodes in the management of epilepsy. Clinical Neurophysiology, 119(1):11–28, 2008.

[31] Farzad Asgarian and Amir M. Sodagar. Wireless telemetry for implantable biomedical microsystems. Biomedical Engineering, Trends in Electronics, Communications and Software, pages 21–44, 2011.

[32] Moo Sung Chae, Zhi Yang, Mehmet R Yuce, Linh Hoang, and Wentai Liu. A 128-channel 6 mw wireless neural recording ic with spike feature extraction and uwb transmitter. Neural Systems and Rehabilitation Engineering, IEEE Transactions on, 17(4):312–321, 2009.

[33] Y.-H. Liu, C.-L. Li, and T.-H. Lin. A 200-pj/b mux-based rf transmitter for implantable multichannel neural recording. IEEE Transactions on Microwave Theory and Techniques, 57(10):2533–2541, 2009.

[34] Mehdi Lotfi Navaii, Mohsen Jalali, and Hamed Sadjedi. An ultra-low power rf interface for wireless-implantable microsystems. Microelectronics Journal, 43(11):848 – 856, 2012.

[35] David A Borton, Ming Yin, Juan Aceros, and Arto Nurmikko. An implantable wireless neural interface for recording cortical circuit dynamics in moving primates. Journal of Neural Engineering, 10(2):026010, 2013.

[36] Helen N. Schwerdt, Wencheng Xu, Sameer Shekhar, Abbas Abbaspour-Tamijani, Bruce C. Towe, Felix A. Miranda, and Junseok Chae. A fully-passive wireless microsystem for recording of neuropotentials using rf backscattering methods. J Microelectromech Syst., 20(5):1119–1130, 2011.

[37] Herve Aubert. Rfid technology for human implant devices. Comptes Rendus Physique, 12.7:675–683, 2011.

[38] Jacopo Olivo, Sandro Carrara, and Giovanni De Micheli. Energy harvesting and remote powering for implantable biosensors. IEEE SENSORS JOURNAL, 11(7):1573, 2011.

[39] B I Rapoport, J T Kedzierski, and R Sarpeshkar. A glucose fuel cell for implantable brain-machine interfaces. PLoS ONE, 7(6):e38436, 2012.

[40] John S. Ho, Alexander J. Yeh, Evgenios Neofytou, Sanghoek Kim, Yuji Tanabe, Bhagat Patlolla, Ramin E. Beygui, and Ada S. Y. Poon. Wireless power transfer to deep-tissue microimplants. PNAS, 111(22):7974–7979, 2014.

[41] Jonas Pistor, Janpeter Hoeffmann, Dagmar Peters-Drolshagen, and Steffen Paul. A programmable neural measurement system for spikes and local field potentials. DTIP Aix-en-Provence, 2011.

[42] J. Pistor, J. Hoeffmann, D. Rotermund, E. Tolstosheeva, T. Schellenberg, D. Boll, V. Gordillo-Gonzalez, S. Mandon, D. Peters-Drolshagen, A K Kreiter, M. Schneider, W. Lang, K. R. Pawelzik, and S. Paul. Development of a fully implantable recording system for ecog signals. Design, Automation and Test in Europe, 2013.

[43] E Tolstosheeva, J Hoeffmann, J Pistor, D Rotermund, T Schellenberg, D Boll, T Hertzberg, V Gordillo-Gonzalez, S Mandon, D Peters-Drolshagen, M Schneider, K R Pawelzik, A K Kreiter, S Paul, and W Lang. Towards a wireless and fully-implantable ecog system. Transducers - The 17th International Conference on Solid-State Sensors, Actuators and Microsystems, page M3P.095, 2013.

[44] E. Tolstosheeva, V. Gordillo-Gonzalez, T. Hertzberg, L. Kempen, I. Michels, A. Kreiter, and W. Lang. A novel flex-rigid and soft-release ecog array. 33rd Annual International IEEE EMBS Conference, 2011.

[45] TI. bqTESLA Portfolio of Wireless Power Solutions. Texas Instruments, 2011.

[46] TI. bq25046EVM-687 Evaluation Module. Texas Instruments, December 2010. SLVU420.

[47] TI. bq51013-Integrated Wireless Power Supply Receiver, Qi (Wireless Power Consortium) Compliant. Texas Instruments, August 2012. SLVSAT9D.

[48] Torex. XCL206-Inductor Built-in Step-Down micro DC/DC Converters. Torex, 2011.

[49] Microsemi. ZL70102-Medical Implantable RF Transceiver MICS RF Telemetry. Microsemi, 2010.

[50] NDK. NZ2016S Series - Crystal Clock Oscillator. NDK, 2013.

[51] J. Hoeffmann, J. Pistor, D. Peters-Drolshagen, E. Tolstosheeva, W. Lang, and St. Paul. Biomedical-asic with reconfigurable data path for in vivo multi/micro-electrode recordings of bio-potentials. MEA Meeting 2012, 2012.

[52] RFMonolithics. RF2607D-403.5 MHz SAW Filter. RF Monolithics, Inc., 2010.

[53] Microsemi. ZL70120 MICS-Band RF Base Station Module (BSM). Microsemi, 2013. Rev. 4.

[54] OrangeTreeTechnologies. ZestET1: GigE TOE & FPGA Module. Orange Tree Technologies, 2013.

[55] Wireless Power Consortium WPC. Wpc to demo worlds most advanced resonant wireless charging system compatible with existing 40+ million qi phones. 2014.

[56] IEEE. Ieee standard for safety levels with respect to human exposure to radio frequency electromagnetic fields, 3 khz to 300 ghz. IEEE Standard for Safety Levels with Respect to Human Exposure to Radio Frequency Electromagnetic Fields, 3 kHz to 300 GHz, page 98, 2006.

[57] G. Charvet, F. Sauter-Starace, M. Foerster, D. Ratel, C. Chabrol, J. Porcherot, S. Robinet, J. Reverdy, R. D’Errico, C. Mestais, and A.L. Benabid. Wimagine: 64-channel ecog recording implant for human applications. Engineering in Medicine and Biology Society (EMBC), 2013 35th Annual International Conference of the IEEE, pages 2756–2759, 2013.

[58] R Muller, Le Hanh-Phuc, Li Wen, P. Ledochowitsch, S. Gambini, T. Bjorninen, A. Koralek, J.M. Carmena, M.M. Maharbiz, E. Alon, and J.M. Rabaey. 24.1 a miniaturized 64-channel 225μw wireless electrocorticographic neural sensor. Solid-State Circuits Conference Digest of Technical Papers (ISSCC), pages 412–412, 2014.

[59] M. Yin, D.A. Borton, J. Aceros, W.R. Patterson, and A. V. Nurmikko. A 100-channel hermetically sealed implantable device for wireless neurosensing applications. Circuits and Systems (ISCAS), pages 2629–2632, 2012.

[60] Masayuki Hirata, Kojiro Matsushita, Takafumi Suzuki, Takeshi Yoshida, Fumihiro Sato, Shayne Morris, Takufumi Yanagisawa, Tetsu Goto, Mitsuo Kawato, and Toshiki Yoshimine. A fully-implantable wireless system for human brain-machine interfaces using brain surface electrodes: W-herbs. IEICE TRANSACTIONS on Communications, E94-B(9):2448–2453, 2011.

[61] J. Aceros, M. Yin, D. Borton, W. Patterson, and A. Nurmikko. A 32-channel fully implantable wireless neurosensor for simultaneous recording from two cortical regions. Annual International Conference of the IEEE Engineering in Medicine and Biology Society, pages 2300–2306, 2011.

[62] Y-K Song, David A Borton, Sunmee Park, William R Patterson, Christopher W Bull, Farah Laiwalla, John Mislow, John D Simeral, John P Donoghue, and Arto V Nurmikko. Active microelectronic neurosensor arrays for implantable brain communication interfaces. IEEE transactions on neural systems and rehabilitation engineering: a publication of the IEEE Engineering in Medicine and Biology Society, 17(4):339, 2009.

[63] Henrique Miranda, Vikash Gilja, Cindy A Chestek, Krishna V Shenoy, and Teresa H Meng. Hermesd: A high-rate long-range wireless transmission system for simultaneous multichannel neural recording applications. IEEE Transactions on Biomedical Circuits and Systems, 4(3):181–191, 2010.

[64] Reid R Harrison, Haleh Fotowat, Raymond Chan, Ryan J Kier, Robert Olberg, Anthony Leonardo, and Fabrizio Gabbiani. Wireless neural/emg telemetry systems for small freely moving animals. IEEE transactions on biomedical circuits and systems, 5(2):103–111, 2011.

[65] Stewart J Thomas, Reid R Harrison, Anthony Leonardo, and Matthew S Reynolds. A battery-free multichannel digital neural/emg telemetry system for flying insects. IEEE transactions on biomedical circuits and systems, 6(5):424–436, 2012.

[66] Hirotaka Sato, Christopher W Berry, Yoav Peeri, Emen Baghoomian, Brendan E Casey, Gabriel Lavella, John M VandenBrooks, Jon Harrison, and Michel M Maharbiz. Remote radio control of insect flight. Frontiers in integrative neuroscience, 3:24, 2009.

[67] A. Sharma, L. Rieth, P. Tathireddy, R. Harrison, H Oppermann, M. Klein, M. Topper, E. Jung, R. Normann, G. Clark, and F. Solzbacher. Evaluation of the packaging and encapsulation reliability in fully integrated, fully wireless 100 channel utah slant electrode array (usea): Implications for long term functionality. 16th International Solid-State Sensors, Actuators and Microsystems Conference, pages 1204–1207, 2011.

[68] Reid R Harrison, Ryan J Kier, Cynthia A Chestek, Vikash Gilja, Paul Nuyujukian, Stephen Ryu, Bradley Greger, Florian Solzbacher, and Krishna V Shenoy. Wireless neural recording with single low-power integrated circuit. IEEE transactions on neural systems and rehabilitation engineering, 17(4):322–329, 2009.

[69] Reid R Harrison, Ryan J Kier, Sohee Kim, Loren Rieth, David J Warren, Noah M Ledbetter, Gregory A Clark, Florian Solzbacher, Cynthia A Chestek, Vikash Gilja, et al. A wireless neural interface for chronic recording. In 2008 IEEE Biomedical Circuits and Systems Conference, pages 125–128. IEEE, 2008.

[70] Sohee Kim, R Bhandari, M Klein, S Negi, L Rieth, P Tathireddy, M Toepper, H Oppermann, and Florian Solzbacher. Integrated wireless neural interface based on the utah electrode array. Biomedical microdevices, 11(2):453–466, 2009.

[71] James C Barrese, Naveen Rao, Kaivon Paroo, Corey Triebwasser, Carlos Vargas-Irwin, Lachlan Franquemont, and John P Donoghue. Failure mode analysis of silicon-based intracortical microelectrode arrays in non-human primates. Journal of Neural Engineering, 10:066014, 2013.

[72] Moo Sung Chae, Zhi Yang, Mehmet R Yuce, Linh Hoang, and Wentai Liu. A 128-channel 6 mw wireless neural recording ic with spike feature extraction and uwb transmitter. IEEE Transactions on Neural Systems and Rehabilitation Engineering, 17(4):312–321, 2009.

[73] Seung Bae Lee, Hyung-Min Lee, Mehdi Kiani, Uei-Ming Jow, and Maysam Ghovanloo. An inductively powered scalable 32-channel wireless neural recording system-on-a-chip for neuroscience applications. IEEE transactions on biomedical circuits and systems, 4(6):360–371, 2010.

[74] Reid R Harrison, Ryan J Kier, Sohee Kim, Loren Rieth, David J Warren, Noah M Ledbetter, Gregory A Clark, Florian Solzbacher, Cynthia A Chestek, Vikash Gilja, et al. 100-channel wireless neural recording system with 54-mb/s data link and 40%-efficiency power link. In Solid State Circuits Conference (A-SSCC), 2012 IEEE Asian, pages 185–188. IEEE, 2012.

[75] Benoit Gosselin. Recent advances in neural recording microsystems. Sensors, 11(5):4572–4597, 2011.

[76] Vikash Gilja, Cindy A Chestek, Ilka Diester, Jaimie M Henderson, Karl Deisseroth, and Krishna V Shenoy. Challenges and opportunities for next-generation intracortically based neural prostheses. IEEE Transactions on Biomedical Engineering, 58(7):1891–1899, 2011.

[77] Kensall D Wise. Wireless integrated microsystems: Wearable and implantable devices for improved health care. In TRANSDUCERS 2009-2009 International Solid-State Sensors, Actuators and Microsystems Conference, pages 1–8. IEEE, 2009.

[78] Reid R Harrison and Cameron Charles. A low-power low-noise cmos amplifier for neural recording applications. IEEE Journal of solid-state circuits, 38(6):958–965, 2003.

[79] Fan Zhang, Jeremy Holleman, and Brian P Otis. Design of ultra-low power biopotential amplifiers for biosignal acquisition applications. IEEE transactions on biomedical circuits and systems, 6(4):344–355, 2012.

[80] Xu Zhang, Weihua Pei, Beiju Huang, Jin Chen, Ning Guan, and Hongda Chen. Implantable microsystem for wireless neural recording applications. In Complex Medical Engineering, 2009. CME. ICME International Conference on, pages 1–4. IEEE, 2009.

[81] Woradorn Wattanapanitch and Rahul Sarpeshkar. A low-power 32-channel digitally programmable neural recording integrated circuit. IEEE Transactions on Biomedical Circuits and Systems, 5(6):592–602, 2011.

[82] Carolina Mora Lopez, Dimiter Prodanov, Dries Braeken, Ivan Gligorijevic, Wolfgang Eberle, Carmen Bartic, Robert Puers, and Georges Gielen. A multichannel integrated circuit for electrical recording of neural activity, with independent channel programmability. IEEE transactions on biomedical circuits and systems, 6(2):101–110, 2012.

[83] A Sharma, L Rieth, P Tathireddy, R Harrison, H Oppermann, M Klein, M Töpper, E Jung, R Normann, G Clark, et al. Evaluation of the packaging and encapsulation reliability in fully integrated, fully wireless 100 channel utah slant electrode array (usea): Implications for long term functionality. Sensors and Actuators A: Physical, 188:167–172, 2012.

[84] Fei Zhang, Mehdi Aghagolzadeh, and Karim Oweiss. A low-power implantable neuroprocessor on nano-fpga for brain machine interface applications. In 2011 IEEE International Conference on Acoustics, Speech and Signal Processing (ICASSP), pages 1593–1596. IEEE, 2011.

[85] John E Ferguson and A David Redish. Wireless communication with implanted medical devices using the conductive properties of the body. Expert review of medical devices, 8(4):427–433, 2011.

[86] Herve Aubert. Rfid technology for human implant devices. Comptes Rendus Physique, 12(7):675–683, 2011.

[87] Farzad Asgarian and Amir M Sodagar. Wireless telemetry for implantable biomedical microsystems. INTECH Open Access Publisher, 2011.

[88] Mehdi Lotfi Navaii, Mohsen Jalali, and Hamed Sadjedi. An ultra-low power rf interface for wireless-implantable microsystems. Microelectronics Journal, 43(11):848–856, 2012.

[89] Mahammad A Hannan, Saad M Abbas, Salina A Samad, and Aini Hussain. Modulation techniques for biomedical implanted devices and their challenges. Sensors, 12(1):297–319, 2011.

[90] Yao-Hong Liu, Cheng-Lung Li, and Tsung-Hsien Lin. A 200-pj/b mux-based rf transmitter for implantable multichannel neural recording. IEEE Transactions on Microwave Theory and Techniques, 57(10):2533–2541, 2009.

[91] Helen N Schwerdt, Wencheng Xu, Sameer Shekhar, Abbas Abbaspour-Tamijani, Bruce C Towe, Félix A Miranda, and Junseok Chae. A fully passive wireless microsystem for recording of neuropotentials using rf backscattering methods. Journal of Microelectromechanical Systems, 20(5):1119–1130, 2011.

[92] Yasaman Damestani, Carissa L Reynolds, Jenny Szu, Mike S Hsu, Yasuhiro Kodera, Devin K Binder, B Hyle Park, Javier E Garay, Masaru P Rao, and Guillermo Aguilar. Transparent nanocrystalline yttria-stabilized-zirconia calvarium prosthesis. Nanomedicine: Nanotechnology, Biology and Medicine, 9(8):1135–1138, 2013.

[93] Jacopo Olivo, Sandro Carrara, and Giovanni De Micheli. Energy harvesting and remote powering for implantable biosensors. IEEE Sensors Journal, 11(EPFL-ARTICLE-152140):1573–1586, 2011.

[94] Benjamin I Rapoport, Jakub T Kedzierski, and Rahul Sarpeshkar. A glucose fuel cell for implantable brain-machine interfaces. PloS one, 7(6):e38436, 2012.

[95] Jonas Pistor, Nils Heidmann, Janpeter Hoffmann, and Steffen Paul. Programmable current source for implantable neural stimulation systems. Procedia Engineering, 87:324–327, 2014.

[96] Dmitry Osipov, Steffen Paul, Serge Strokov, and Andreas K Kreiter. A new current stimulator architecture for visual cortex stimulation. In Nordic Circuits and Systems Conference (NORCAS): NORCHIP & International Symposium on System-on-Chip (SoC), 2015, pages 1–4. IEEE, 2015.

[97] Dmitry Osipov and Steffen Paul. A novel hv-switch scheme with gate-source over-voltage protection for bidirectional neural interfaces. In 2015 IEEE International Conference on Electronics, Circuits, and Systems (ICECS), pages 25–28. IEEE, 2015.

